# All-optical mechanobiology interrogation of YAP in human cancer and normal cells using a novel multi-functional system

**DOI:** 10.1101/2021.06.09.447829

**Authors:** Qin Luo, Miao Huang, Chenyu Liang, Justin Zhang, Gaoming Lin, Sydney Yu, Mai Tanaka, Sharon Lepler, Juan Guan, Dietmar Siemann, Xin Tang

## Abstract

Long-term multi-functional imaging and analysis of live cells require streamlined functional coordination of various hardware and software platforms. However, manual control of various equipment produced by different manufacturers is labor-intensive and time-consuming, potentially decreasing the accuracy, reproducibility, and quality of acquired data. Therefore, an all-in-one and user-programmable system that enables automatic, multi-functional, and long-term image acquisition and is compatible with most fluorescent microscopy platforms can desirably benefit the scientific community. In this paper, we introduce the full operating protocols of utilizing a novel integrated software system that consists of (1) a home-built software program, titled “Automatic Multi-functional Integration Program (AMFIP)”, which enables automatic multi-channel imaging acquisition, and (2) a suite of quantitative imaging analysis and cell traction computation packages. We applied this integrated system to reveal the previously unknown relationship between the spatial-temporal distribution of mechano-sensitive Yes-associated protein (YAP) and the cell mechanics, including cell spreading and traction, in human normal cells (B2B) and lung cancer cells (PC9). Leveraging our system’s capability of multi-channel control and readout, we found: (1) B2B normal cells and PC9 cancer cells show distinct YAP expression, traction, and cell dynamics relationship during cell spreading and migration processes; and (2) PC9 cancer cells apply noticeable peri-nuclear forces on substrates. In summary, this paper presents a detailed stepwise protocol on how to utilize an integrated user-programmable system that enables automatic multi-functional imaging and analysis to elucidate YAP mechanosensitivity. These tools open the possibility for detailed explorations of multifaceted signaling dynamics in the context of cell physiology and pathology.

**SUMMARY:** This paper presents a detailed stepwise protocol on how to utilize an integrated user-programmable system that enables all-optical electro-mechanobiology interrogation to elucidate YAP mechanosensitivity.

## INTRODUCTION

An all-in-one imaging program that enables automatic coordination of multi-functional optoelectronic devices will reduce labor-intensive and error-prone manual operations and is essential for researchers to conduct long-term live-cell imaging [1]–[4]. However, most existing public programs in the biomedical research community either only apply to limited optoelectronic devices or require additional hardware for the coordination of different equipment. We recently developed an open-source and software-based program, titled “Automatic Multi-functional Integration Program (AMFIP)”, that enables multi-channel and time-lapse imaging [5]. Developed as a plugin of µManager [6], [7], AMFIP executes customized Java scripts to accomplish software-based communications of multiple optoelectronic hardware and software platforms, including but not limited to those from Nikon. The utilization of AMFIP avoids the demand of purchasing additional controlling devices that may not be available for and compatible with every imaging system.

In this protocol paper, we provide an integrated experimental and computational system that combines AMFIP with digital imaging analysis and cell traction force microscopy to elucidate the distinct YAP mechanobiology in human normal B2B (**Figure 1**) and lung cancer PC9 (**Figure 2**) cell lines. The protocol presents how to (1) apply AMFIP to conduct automatic long-term imaging for both cell lines that express mNEonGreen2-tagged YAP; and (2) combine Fiji ImageJ, MATLAB, and Origin for the quantitative analysis of YAP nuclear/cytoplasm (N/C) ratio based on their fluorescent intensity (**Figures 3 and 4**), cellular displacement field (**Figures 1B and 2B**), and cellular traction field (**Figures 1C and 2C**). The results suggest that (1) during the first 10 hours of cell spreading, the YAP N/C ratio of single B2B cells shows more evident time-dependent variation and fluctuation compared to that of single PC9 cells (**Figures 5 and 6**); and (2) PC9 cancer cells generate noticeable traction in their peri-nuclear regions (**Figure 7**).

**Figure 1.**
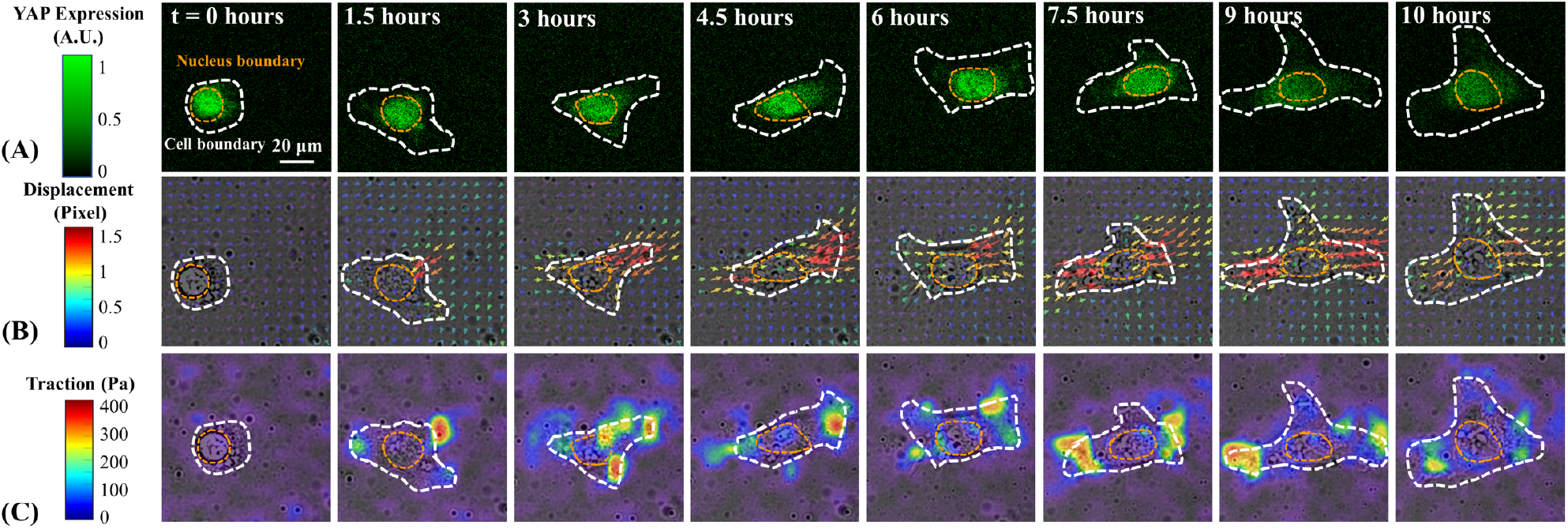
The changes in YAP expression/distribution, substrate displacement field, and traction field of a B2B normal cell during early spreading. The B2B cell was seeded on 5-kPa PAA gel and imaged for over 10 hours after attaching. (**A**) YAP expression is represented by green fluorescence intensity. Note: The YAP intensity inside the nucleus gradually decreases but remains higher over time. **(B)** Substrate deformation (overlapped with the bright-field image) at cell location is represented by displacement field at each time point. Displacement direction and magnitude are shown by the arrow direction and color, respectively. The displacement becomes larger at the ends of the B2B cell body as the cell spread area increases. **(C)** Traction field (overlapped with the bright-field image) calculated from displacement field. The traction is concentrated on the boundary of B2B cells.

**Figure 2.**
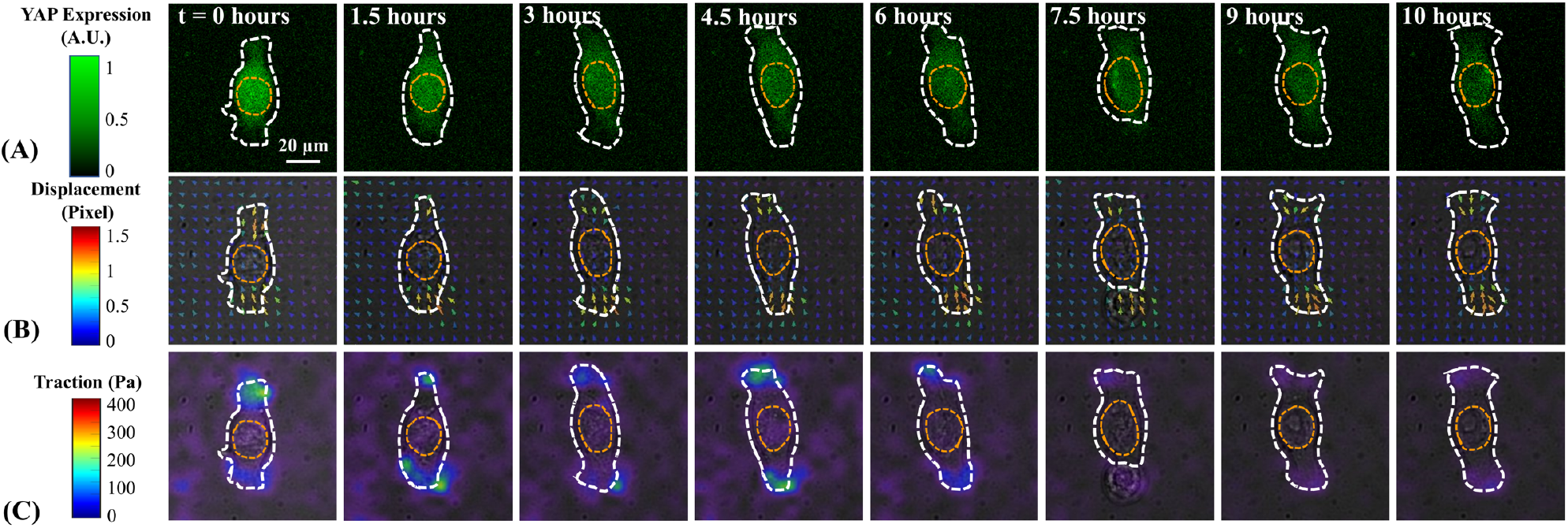
The changes in YAP expression/distribution, substrate displacement field, and traction field of a PC9 cancer cell during early spreading. The PC9 cells was seeded on 5-kPa PAA gel and imaged for over 10 hours after attaching. (**A**) YAP expression is represented by green fluorescence intensity. Note: The YAP intensity becomes homogenous from the 1.5^th^ hour on. **(B)** Substrate deformation (overlapped with the bright-field image) at cell location is represented by fluorescent beads displacement field at each time point. Displacement direction and magnitude are shown by the arrow direction and color, respectively. The displacement field caused by PC9 cells is smaller compared to that caused by the B2B cell. Throughout the 10-hour spreading process, the area of PC9 cells nearly remains constant. **(C)** Traction field (overlapped with the bright-field image) calculated from displacement field. The traction generated by this representative PC9 cell gradually decreases from the 6^th^ hour to the 10^th^ hour.

**Figure 3.**
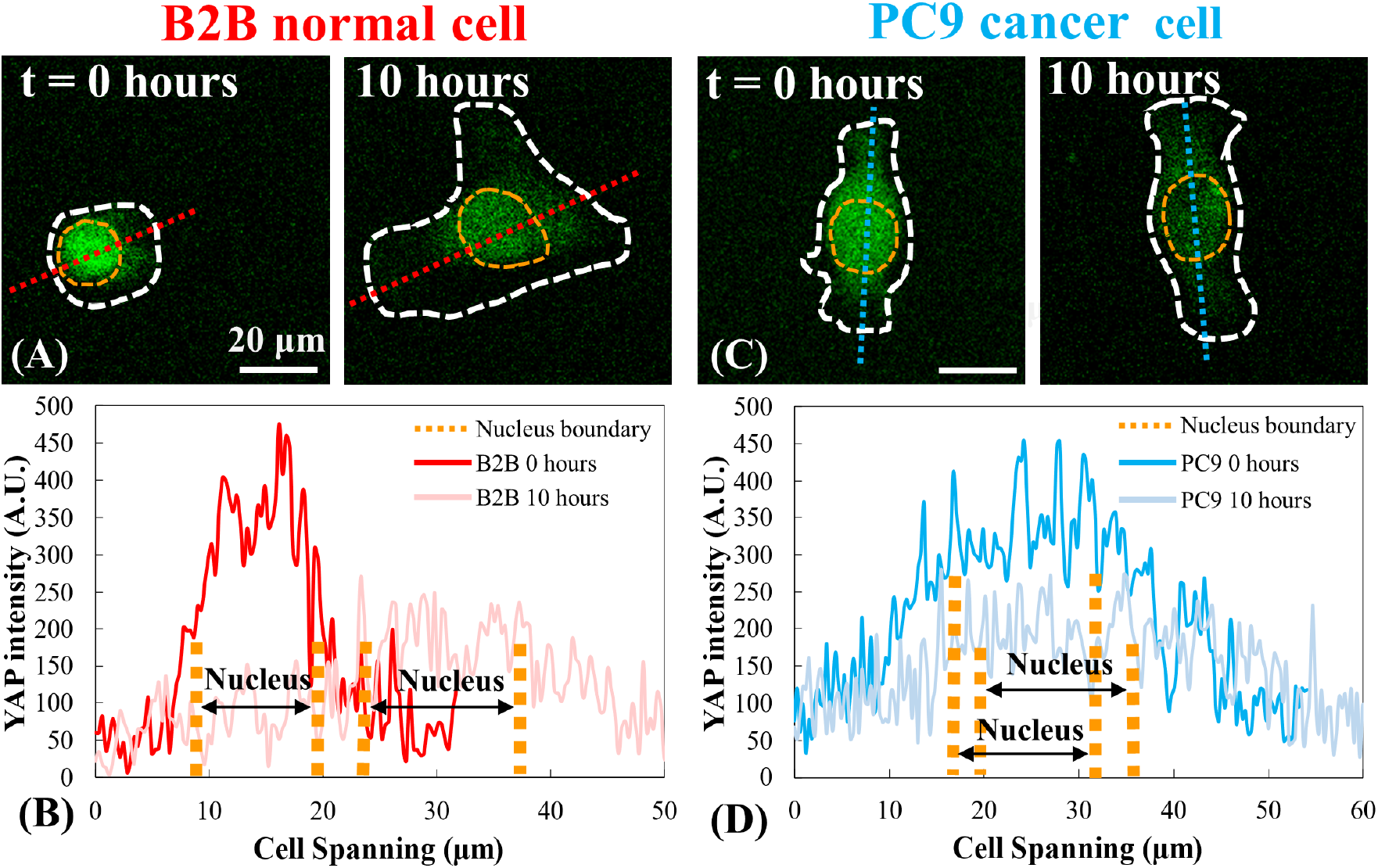
YAP distribution in B2B and PC9 cells at early spread stage. **(A)** YAP intensity of the B2B cell is measured along the assigned red axis at the 0^th^ and the 10^th^ hour. **(B)** At the 0^th^ hour, YAP intensity shows dramatic concentration differences between nucleus and cytoplasm. At 10^th^ hour, YAP intensity becomes more homogenous across the whole cell body. **(C)** YAP intensity of the PC9 cell is measured along the assigned blue axis at the 0^th^ and the 10^th^ hour. **(D)** At the 0^th^ hour, YAP intensity in the nucleus appears higher than that in the cytoplasm, but the distinction is not as evident as that in B2B cells. At the 10^th^ hour, YAP intensity in the nucleus still appears slightly higher than that in the cytoplasm, but the trend of variation is similar with that at the 0^th^ hour.

**Figure 4.**
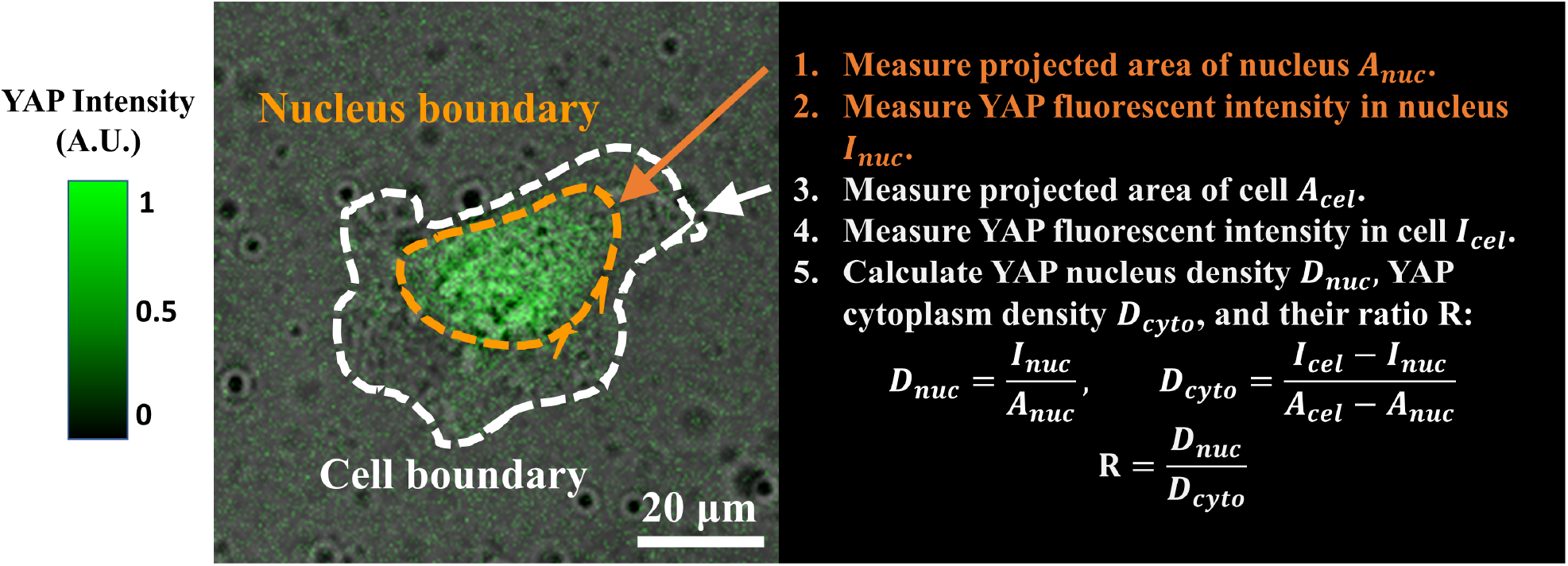
The procedure of measuring YAP N/C ratio. (1) Apply Fiji ImageJ to draw the outline of nucleus and measure its 2D projected area *A*_*nuc*_. (2) Measure the intensity of fluorescence inside the nucleus *I*_*nuc*_. (3) Draw the outline of cell body and measure its projected area *A*_*cel*_. (4) Measure the Measure the intensity of fluorescence inside the cell *I*_*cel*_. (5) Calculate YAP nucleus density *D*_*nuc*_, YAP cytoplasm density *D*_*cyto*_, and their ratio *R*: *D*_*nuc*_=*I*_*nuc*_/*A*_*nuc*_; *D*_*cyto*_=(*I*_*cel*_*−I*_*nuc*_)/(*A*_*cel*_*−A*_*nuc*_); *R*=*D*_*nuc*_/*D*_*cyto*_.

**Figure 5.**
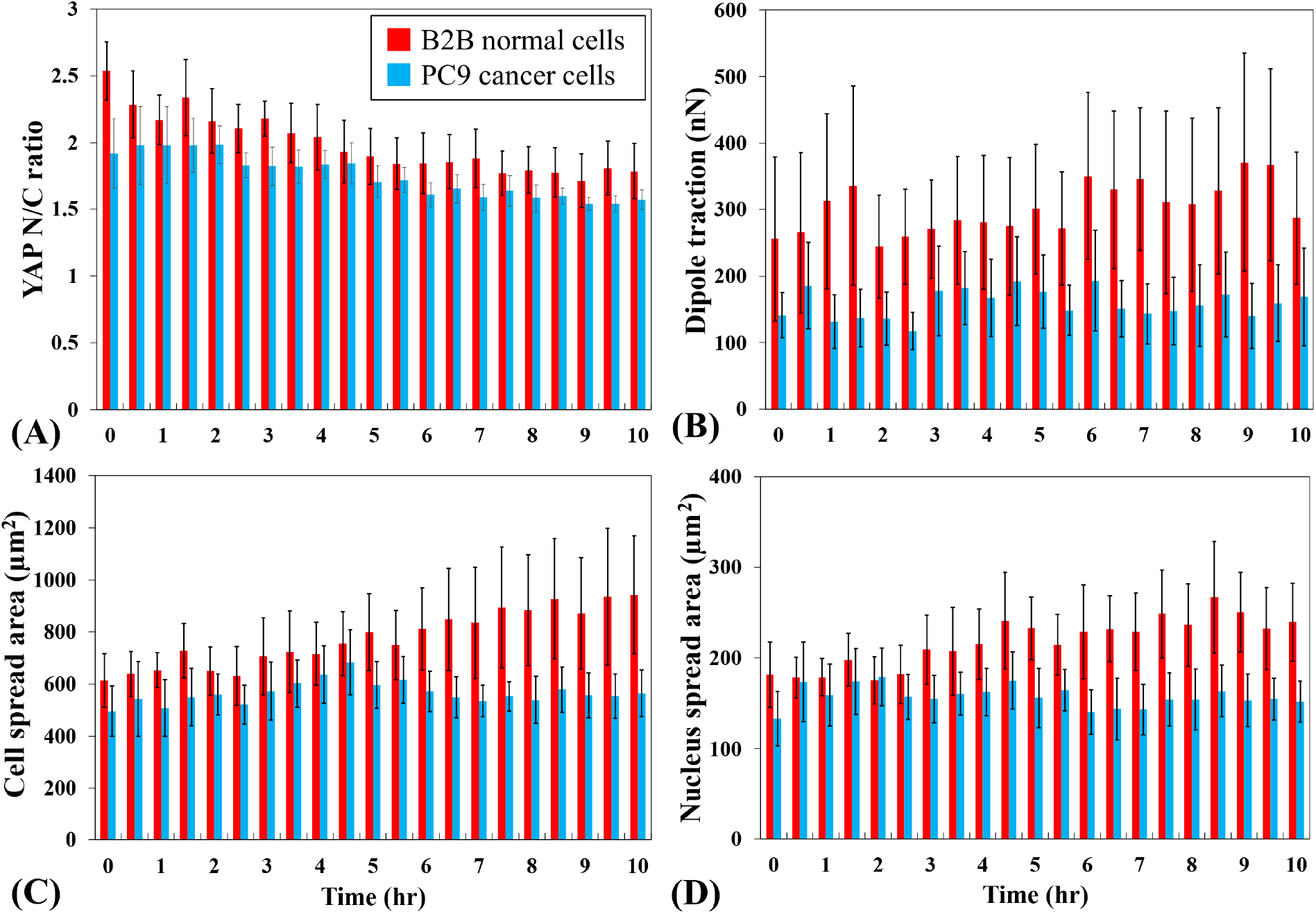
Distinct YAP expression, cell/nucleus morphology and cellular traction in PC9 cancer and B2B normal cells during cell spreading. **(A)** YAP N/C ratio change during the first 10 hours of single cell spreading. Average YAP N/C ratio of B2B cells (red column; n = 10) changed from 2.54 ± 0.22 to 1.79 ± 0.21 (n = 10; p-value = 0.0022**) while the average YAP N/C ratio of PC9 cells (blue column; n= 5) changed from 1.92 ± 0.26 to 1.57 ± 0.07 (p-value = 0.187 (not significant (ns))). **(B)** The average dipole traction as a function of time. The average dipole traction of B2B cells changed from 256.17 ± 123.69 nN to 287.44 ± 99.79 nN (p-value = 0.7593 (ns)) and the average dipole traction of PC9 cells changed from 141.19 ± 33.62 nN to 168.52 ± 73.01 nN (p-value = 0.7137 (ns)). **(C)** The average cell area as a function of time. The average cell spread area of B2B cells increased from 613.89 ± 102.43 µm^2^ to 942.51 ± 226.71 µm^2^ (p-value = 0.0512 (ns)) and the average cell spread area of PC9 cells changed from 495.78 ± 97.04 µm^2^ to 563.95 ± 89.92 µm^2^ (p-value = 0.5804 (ns)). **(D)** The average nucleus area as a function of time. The average nucleus spread area of B2B cells increased from 181.55 ± 36.18 µm^2^ to 239.38 ± 43.12 µm^2^ (p-value = 0.1217 (ns)) and the average nucleus spread area of PC9 cells changed from 133.31 ± 30.05 µm^2^ to 151.93 ± 22.49 µm^2^ (p-value = 0.5944 (ns)).

**Figure 6.**
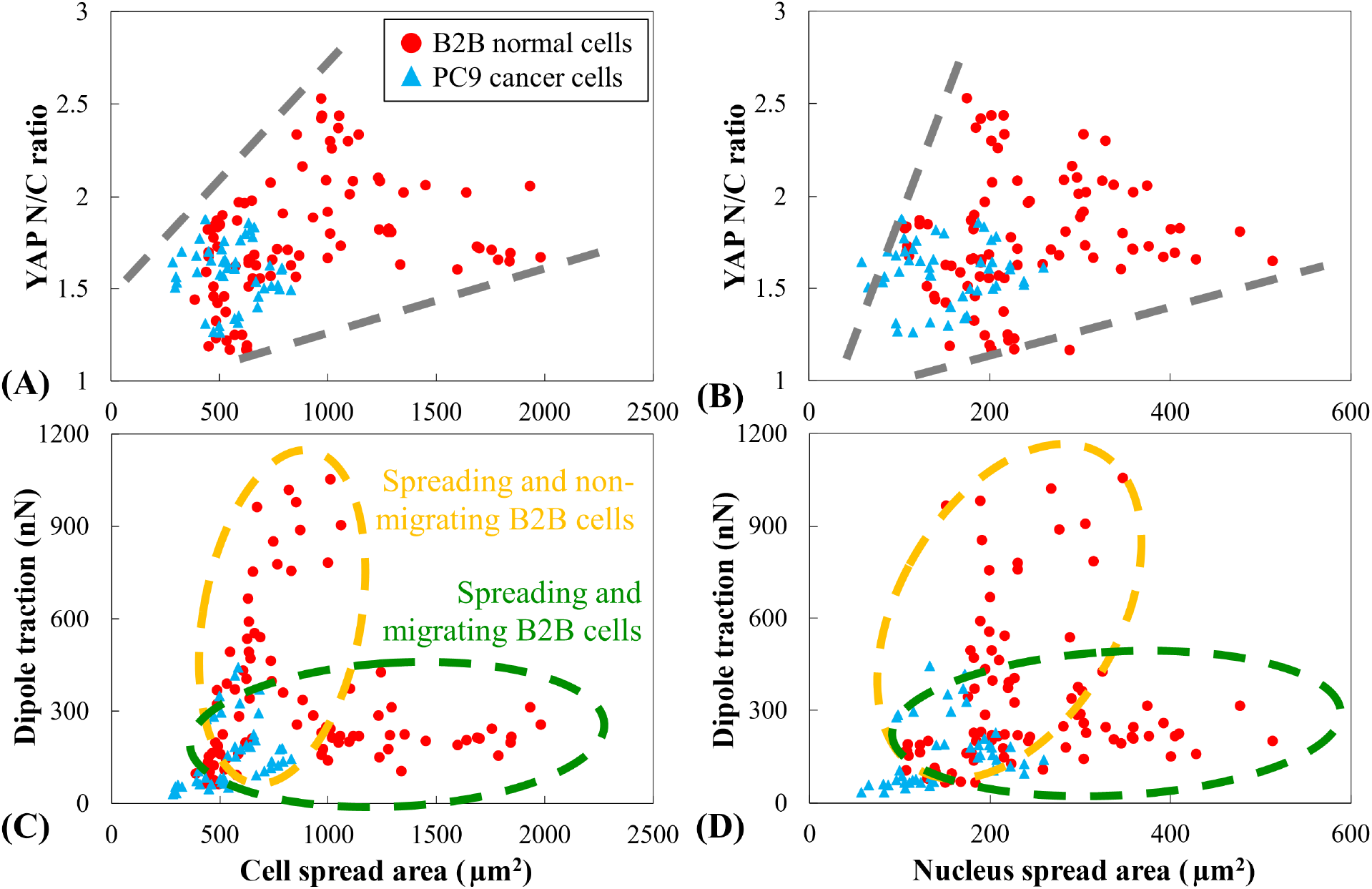
YAP N/C ratio and dipole traction force as a function of spread areas of cell and nucleus. YAP N/C ratio and dipole traction of B2B cells (n=10) and PC9 cells (n=5) are calculated from the 6^th^ hour to the 10^th^ hour after attaching to substrate. **(A)** YAP N/C ratio as a function of cell spread area. The YAP N/C ratios of B2B cells vary from 1.16 to 2.53, while the YAP N/C ratios of PC9 cells vary from 1.27 to 1.88. The cell spread areas of B2B cells vary from 391.94 µm^2^ to 1986.40 µm^2^. The cell spread area of PC9 cells range from 284.46 µm^2^ to 830.12 µm^2^. **(B)** YAP N/C ratio as a function of nucleus spread area. The nucleus spread areas of B2B cells vary from 107.09 µm^2^ to 514.28 µm^2^. The nucleus spread areas of PC9 cells range from 58.03 µm^2^ to 259.65 µm^2^. Dipole traction of B2B cells as a function of cell spread area (**C**) and nucleus spread area (**D**). Spreading and non-migrating B2B cells show higher traction (from 47.50 nN to 1051.48 nN) with lower cell and nucleus area. While spreading and migrating, B2B cells show lower traction (from 105.80 nN to 310.28 nN) with larger ranges of cell and nucleus areas.

**Figure 7.**
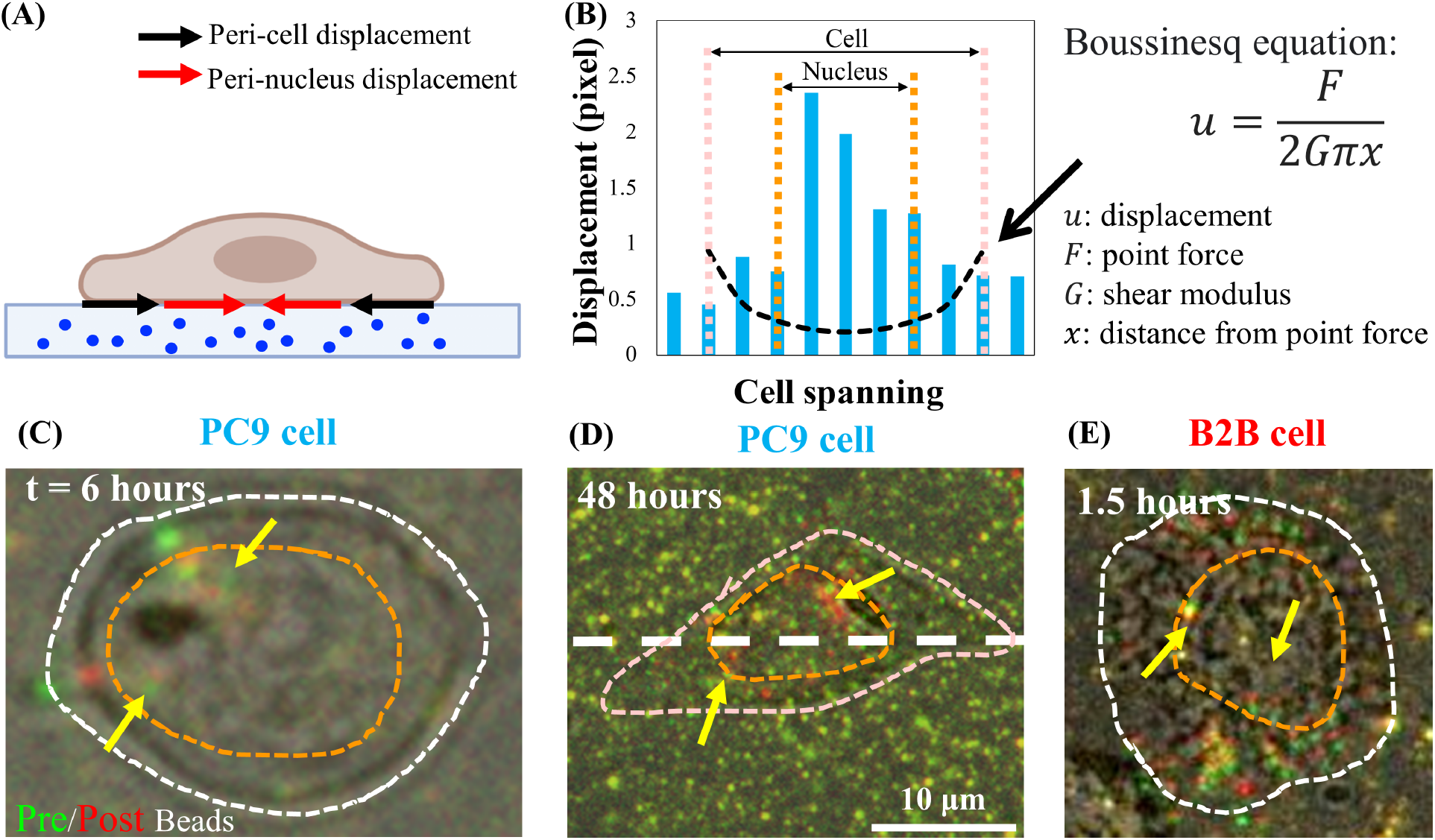
Peri-nuclear displacement in normal B2B and cancer PC9 cells. **(A)** Schematic side-view diagram of peri-nucleus and peri-cell displacement measured from beads displacement in the substrate. **(B)** Substrate displacement underneath the PC9 cell is measured along the cell axis (white dashed line in **7D**). The theoretical displacement generated by dipole force at cell boundary is shown by the Boussinesq equation (black dashed line). **(C) ̲ (D)** Overlapped fluorescent beads images with (red) and without (green) cell for the PC9 cells at the 6^th^ hours after attaching (Top view). Yellow (totally merged red and green) beads indicate no displacement. The separated green and red beads (pointed by yellow arrows) represent the peri-nucleus displacement. Yellow arrows indicate these contracted per-nucleus spots. **(E)** Peri-nuclear displacement generated by B2B cell at 1.5^th^ hour after the cell-substrate attachment.

The integrated system and methodologies described in this protocol transcend the specific cell types. Researchers can apply the protocol to customize their specific live-cell imaging experiments and elucidate multifaceted signaling dynamics in the context of cell physiology and pathology (**Figure 8**). We have established a new all-optical electrophysiology (Optopatch) interrogation theme to explore the previously inaccessible bioelectricity of cancer cells in our laboratory.

**Figure 8:**
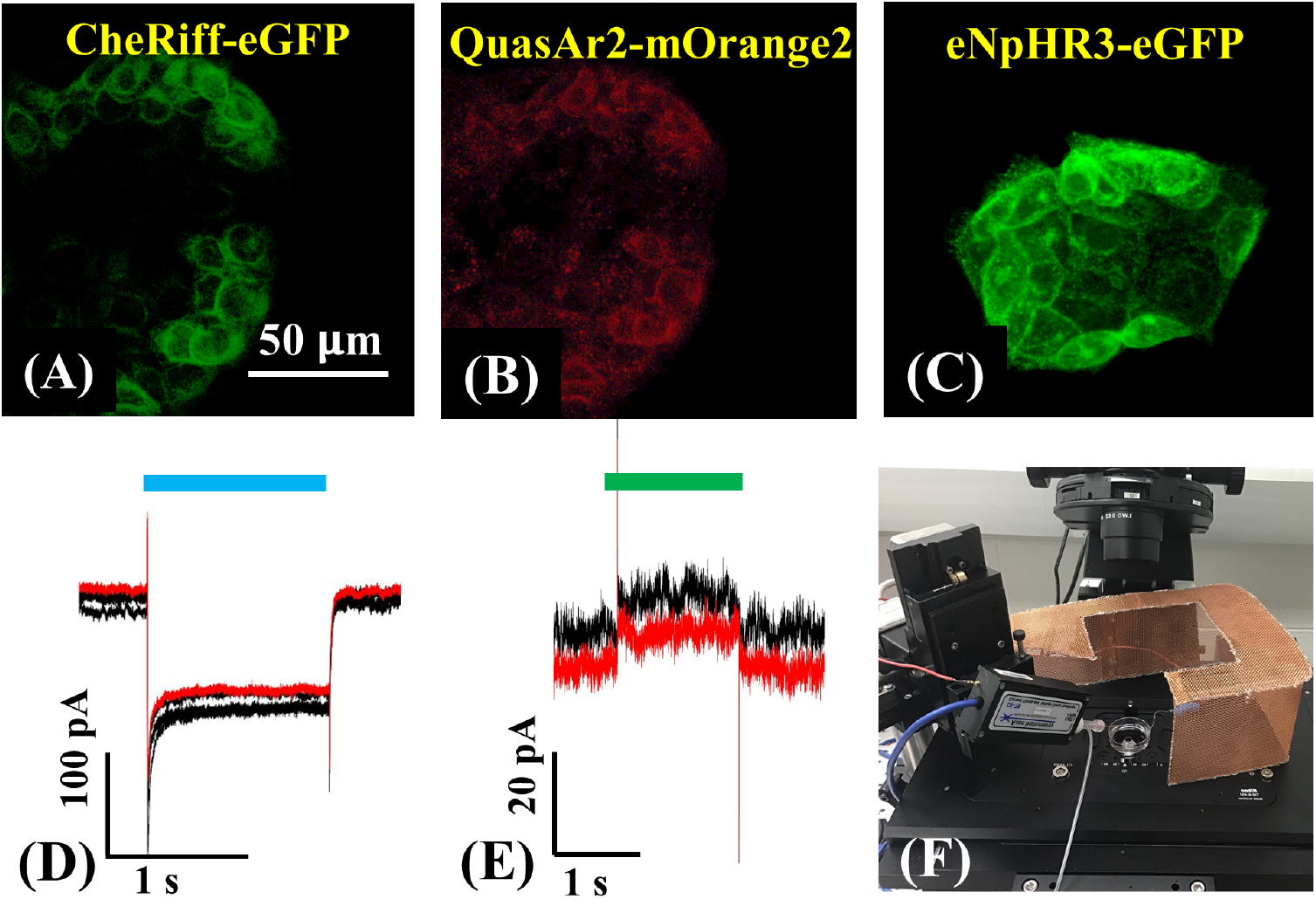
All-optical electrophysiology (Optopatch) interrogation in human cancer cells. Human colon cancer cells HCT8 expressing **(A)** CheRiff; **(B)** QuasAr2; and **(C)** eNpHR3. Blue and green laser lines stimulated the membrane depolarization **(D)** and hyperpolarization of HCT-8 cells. **(E)** Trans-membrane currents are recorded by a manual patch clamp. **(F)** Electrophysiology rig and Nikon A1R fluorescent confocal microscope on which the all-optical electrophysiological control and readout are conducted.

## PROTOCOL

### 1. Generation of endogenously tagged mNeonGreen21-10/11 cell lines: the human lung cancer cell line (PC9) and the human bronchial epithelial cell line (Beas2B)

1. Insert the DNA sequence coding the 11th strand of fluorescence protein mNeonGreen2 into the YAP genomic locus through the CRISPR-Cas9 gene-editing system. The detailed molecular biology protocols are reported elsewhere [8].
2. Check the mNeonGreen2 expression using an epifluorescence microscope. mNeonGreen2 is tagged to YAP whenever the cell expresses YAP in the context of its native gene regulatory network.
3. Confirm the correct integration of mNeonGreen211 by genomic sequencing and by the reduction in fluorescence upon gene knockdown.
4. Collect the cells with the tagged protein of interest through fluorescence-activated cell sorting (FACS).

### 2. Maintenance of PC9 and B2B cells

1. Maintain both cell lines in humidified incubators with 5 % CO_2_ at 37 °C.
2. Culture endogenously tagged PC9 and Beas2B cells in RPMI-1640 medium supplemented with 10 % FBS and penicillin-streptomycin at 100 µg/mL.
3. Test all cell lines for mycoplasma every 3 months using MycoAlert Mycoplasma Detection Kit (Lonzha).
4. Use the cell lines (<20 passages) from thaw for all the experiments described in this protocol.

### 3. Setup of hardware and software environment

1. Hardware environment setup of the experiment
  1. Connect the Nikon A1R confocal controller and the Nikon Ti2-E inverted microscope to the computer.
  2. Install the Nikon NIS-Elements (Elements) software platform.
  3. Turn on the Nikon A1R confocal controller and the Ti2-E microscope. Next, launch Elements.
  4. Open the control panels of confocal, laser, and Ti2-E in Elements. Next, check whether the three panels function properly.
2. Software environment setup of AMFIP
  1. Install IntelliJ (JetBrains), Java Development Kit 14.0, µManager version 2.0 gamma, and Fiji ImageJ (National Institutes of Health and the Laboratory for Optical and Computational Instrumentation) on the computer.
  2. Open the AMFIP project downloaded from GitHub (link: https://github.com/njheadshotz/AMFIP) in IntelliJ.
  3. Click **Settings > Compiler > Annotation Processors** and check **Enable annotation processing**.
  4. Click **Project Structure > Artifacts** and create a JAR file. Set the output directory to **mmplugins** under the directory of µManager.
  5. Click **Project Structure > Libraries** and add **mmplugins** and **plugins** under the directory of µManager.
  6. Click **add Configuration** under the **Run** drop-down menu and create an application.
  7. Enter **ij.ImageJ** into the **Main class**.
  8. Enter **-Xmx3000m - Dforce.annotation.index=true** into **VM option**.
  9. Set the directory of µManager to the **Work directory**.
  10. Click **Run** to activate µManager with the AMFIP plugin.
3. Connect µManager with the Nikon Ti2-E microscope
  1. Add the adaptive driver of the Nikon Ti2-E microscope (Link: https://micro-manager.org/wiki/NikonTi2) into the directory of µManager.
  2. Open µManager. Click **Devices >Hardware Configuration Wizard** and create a new configuration.
  3. Add the Nikon Ti2 driver under **Available Devices**.
  4. Select all peripheral devices and save the new configuration file.
  5. Restart µManager and select the configuration file in step 4 in **Micro-Manager Startup Configuration**.

### 4. Gel preparation

Polyacrylamide (PAA) hydrogel substrates (with 2 % of fluorescent beads (Invitrogen)) that have Elastic Modulus 2 kPa, 5 kPa, 10 kPa, and 40 kPa are prepared following the established protocols [9], [10].

### 5. Cell culture

1. Bond the glass coverslips with PAA hydrogels to the glass-bottom petri dish (MatTek) to avoid physical drift of gels during cell seeding and imaging processes
  1. Lift the coverslip (with the PAA hydrogel on top) with a sterilized clean tweezer out from the petri dish that was used to contain the prepared gels.
  2. Apply a dry Kimwipe (Kimtech Science) to clean up water droplets at the bottom surface of the glass coverslip.
  3. Apply the sterilized tweezer to hold the glass coverslip.
  4. Drip a small droplet (1−5 µL) of super glue (Gorilla) at the two diagonal corners on the bottom side.
  5. Apply sterilized Kimwipe to remove extra droplets of super glue.
  6. Apply the sterilized tweezer to restore the coverslip into the glass-bottom petri dish. Slightly press the corner of the coverslip to ensure the droplets of glue have full contact with the surface of the petri dish.
  7. Place the lid back onto the petri dish to minimize the evaporation of PBS in the PAA hydrogels. Wait for 3 minutes to allow the super glue to solidify and dry in the petri dish.
  8. Fill PBS (4 mL) into the petri dish.
  9. Repeat the above steps 1−8 to the rest of the PAA hydrogel samples in the petri dishes used for imaging.
  10. Apply 75 % ethanol to sterilize the outer surface of all petri dishes prepared and put it into the tissue culture bio-hood. Turn on the ultraviolet light for 5 minutes and sterilize the samples.
2. Seed the cells onto the gel top surface
  1. Turn off the ultraviolet light. Next, spray 75 % ethanol to the rubber gloves on hands (repeat this process every time before opening the incubator) and take out the flask (with B2B/PC9 cells inside it) from the incubator (37 °C) into the bio-hood. Use an aspirating pipette connected to a vacuum pump to suck out all the culture medium (Gibco; 11875-093) and add 5 mL PBS to wash the flask.
  2. Add 2 mL of 0.05 % Trypsin (Corning) to detach the cells from the bottom of the flask.
  3. Place the flask in the incubator. Wait for 5 minutes.
  4. Take out the flask to the bio-hood. Add 8 mL new culture medium into the flask and pipette the solution for several times.
  5. Completely transfer the 10 mL solution to a small tube. Centrifuge the tube with another tube containing 10 mL water (acceleration: 300 G) for 5 minutes.
  6. Check the cell pellet at the bottom of the tube. Slowly flatten the tube and use the aspirating pipette to suck out all the culture medium in the tube (avoid touching the cell pellet). Next, add 8 mL of new culture medium and pipette the solution several times until all cells are mixed with the medium.
  7. Deposit 100 µL of the medium with cells onto the gel surface. Wait for 5 minutes. Next, slowly add 4 mL of new culture medium into the petri dishes (avoid directly dropping the new medium onto the gel).
  8. Place the petri dish into the incubator (37 °C). Wait to allow cells to attach to the gel surface (B2B: 0.5-1 hour; PC9: 4-5 hours).

### 6. Cell Imaging

AMFIP realizes automatic, multi-channel, and long-term imaging by coordinating with different hardware and software systems: (1) AMFIP manipulates µManager to automatically move the motor-stage of the Ti2-E microscope to multiple field-of-views (FOVs) and acquire bright-field images through a monochrome camera (BFS-U3-70S7M-C; FLIR); and (2) AMFIP activates multiple macro files inside Elements with a customized Java script to accomplish automatic operations for confocal z-stack imaging and the switch of different laser channels (405 nm and 488 nm).

1. Set the environment for long-term imaging
  1. Place the environment chamber (Tokai Hit) onto the motor-stage of the Ti2-E microscope. Set the CO2 flow rate to 160 and adjust the temperature of the chamber (Top: 44 °C; Bath: 42 °C; Stage: 40 °C). Next, add purified water into the bath of the chamber.
  2. Take out the glass-bottom petri dish with cells from the incubator and place into the chamber.
  3. Turn on the A1R confocal controller and the Ti2-E microscope. Switch the light path to the right and observe the cell attaching using SpinView or µManager.
  4. If sufficient cells have attached to the gel, put the petri-dish back in the incubator.
  5. Cut two small pieces of Scotch tape and stick them to the chamber around the circular hole. Next, put very little super glue onto the tape (only at the area that will be covered by the petri dish).
  6. Take out the petri dish from the incubator. Next, slowly put down the petri dish to the chamber and let the bottom of the petri dish make contact with the dropped super glue.
  7. Press the lid of the petri dish for 1 minute to allow the super glue to make full contact with the petri-dish and solidify. Next, gently push the petri dish horizontally to confirm that the petri dish is unmovable on the chamber.
  8. Close the lid of the chamber.
2. Set the image acquisition for bright-field images
  1. Open **IntelliJ** and set a parameter: T1 in Elements_script. java file (line 93). This value must be larger than the running time of the macro in Elements used for the confocal imaging.
  2. Run the **AMFIP IntelliJ project**.
  3. Select **Live** and **Multi-D Acq**. button on the main interface of µManager. Next, switch the light path of the Ti2-E microscope to the right for bright-field imaging, switch to 10× objective, and open Dialamp (the light source for bright-field imaging; intensity: 5 %).
  4. Adjust the XY joystick and the knob of the Z-plane to find the correct position of gel on the petri dish.
  5. Use a 10× objective to find appropriate FOVs of multiple single cells attached to the gel.
  6. Click the **Multiple Positions (XY)** button on the **Multi-Dimensional Acquisition** window. The **Stage Position Lis**t window pops up. Next, change the objective to 40×, increase the intensity of DiaLamp to 15 %, re-adjust the XY-motor-stage to locate the FOVs, and record the coordinates by clicking the **Mark** button on the Stage Position List window.
  7. Record 6-7 FOVs desired. Click the **Save As**… button on the Stage Position List window to save the recorded coordinates. Next, set the time interval of imaging acquisition to T1.
3. Set the image acquisition for 2D-YAP and beads images
  1. Open **Elements**, change the light path to the right for confocal imaging, and turn off the DiaLamp. Next, click the **Remove Interlock** button and turn on the **FITC laser channel** (for YAP imaging).
  2. Adjust the **second per frame** to 1/2 and spin the knob of the Z-plane to quickly find the Z-position of the attached cells. Record the lowest position and the highest position.
  3. Select **Macro Editor** under the **Macro** drop-down menu and input the values from step 2 to a macro file.
  4. Turn on the **DAPI laser channel** (for beads imaging) to find and record the focused Z-position of beads.
  5. Go to **Macro editor** and input the recorded values into the macro file.
4. Set the task of moving the motor-stage using AMFIP
  1. Go to **µManager** and click **Plugins > Automation** to open the GUI of AMFIP.
  2. Click **Add Point** or **Remove point** buttons to acquire the exact number of FOVs selected. Input the recorded coordinates of FOVs into the **Coordinates Panel**.
  3. Define the total experiment time in the **Total Experiment Time** text field.
  4. Click the **Additional Time Configuration** button and define the time interval of moving the motor-stage to each FOV.
  5. Maximize the window size of Elements and drag the GUI of AMFIP to the right side of the screen (avoid the GUI of AMFIP disturbing the automatic operations of the cursor).
  6. Click **Enter**.
5. Dissolve the cells after the image acquisition
  1. After finishing the long-term imaging, stop the task of AMFIP.
  2. Open **Elements** and set a Z-stack imaging (set the Z-range to be larger than the Z-range of the beads).
  3. Switch the light path to the right and open the DiaLamp (intensity: 15 %).
  4. Slowly and carefully remove the lids of the chamber and the petri dish. Meanwhile, monitor the bright-field view in case any drift of the FOV happens.
  5. Apply a plastic pipette to suck up 0.5 mL sodium dodecyl sulfate (SDS) solution, carefully hold the plastic pipette to be a little above the culture medium and add 1-2 droplets of SDS solution into the culture medium.
  6. Once the cells in the bright-field view are dissolved, switch the light path to the left, close DiaLamp, click the Remove Interlock button.
  7. Run the Z-stack imaging. Save the image stack and name it as “Reference_N” (N is the sequence number of each FOV).
  8. Click the **Multiple Positions (XY)** button on the **Multi-Dimensional Acquisition** window. Next, select the next FOV and click the **go to** button to move the motor-stage to the second FOV.
  9. Repeat step 7 for each FOV.

### 7. Measurement of YAP N/C ratio

Perform image analysis for YAP N/C ratio using the Fiji ImageJ software (**Figure 4**).

1. Open **Fiji ImageJ**. Import the bright-field image stack for all FOVs acquired by µManager.
2. Open the **Image** drop-down menu and select **Stacks > Tools > Slice Keeper**. Next, export the bright-field image stack for each FOV.
3. Import the fluorescence image of the FITC channel and overlay it with the bright-field image at the same FOV. To do this, choose the fluorescent image and select **Overlay > Add Image**… (Image to add: the bright-field image; X and Y location depends on the size of the bright-field image acquired by different cameras; Opacity: 60-70).
4. Open the **Analyze** drop-down menu and select **Set Measurements…**. Select **Area**; **Integrated density** and **Mean gray value**.
5. Click the **Freehand selections** button on the main interface of ImageJ.
6. Draw the outline of the cell body and the nucleus desired. Next, click **Analyze > Measure** or press the M button on the keyboard.
7. The **Results** window pops up. The values under the **Area** column represent the area of the selected region (µm^2^). The values under the **IntDen** column represent the fluorescence intensity of the selected region.
8. Calculate the YAP N/C ratio following the formula:

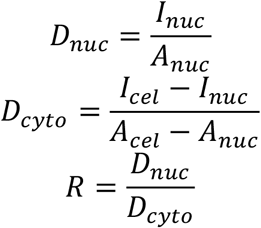 Where *I*_*nuc*_ and *I*_*cel*_ represent the relative intensity of the nucleus and the cell body, and *A*_*nuc*_ and *A*_*cel*_ represent the area of the nucleus and the cell body. *R* is the YAP N/C ratio.
9. Save the outlines for future calculation of dipole traction force and peri-cell/peri-nuclear displacement. To do this, click **Analyze > Tools > Save XY Coordinates**…

### 8. Measurement of traction field

1. Apply traction force microscopy through Fiji ImageJ plugins [11], [12].
  1. Open **Fiji ImageJ**.
  2. Import the image stack of beads for a FOV.
  3. Select the slice that shows the clearest distribution of beads and extract it using **Slice Keeper**.
  4. Import the image stack of the reference at the same FOV.
  5. Choose the slice with the same brightness and contrast as the slice in step 3. Next, extract it as a reference image.
  6. Select **Images > Stacks > Tools > Concatenate** to combine the two slices from steps 3 and 5 (select the reference image as the first slice).
  7. Select **Plugins > Template Matching > Align slices in stack** or **Plugins > Image Stabilizer** to align the two slices.
  8. Select **Image > Stacks > Stack to Images**. Next, select **Image > Lookup Tables > Green** to convert the color of the first slice to green and select **Image > Lookup Tables > Red** to convert the color of the second slice to red.
  9. Select **Image > Color > Merge Channels** to merge the two images.
  10. Overlap the image with the bright-field image at the same FOV. This overlapped image is used to observe beads displacement.
  11. Select **Plugins > PIV > iterativ PIV(Basic)**…. Set the interrogation window size to 128/256; 64/128; 32/64 (at least four beads per interrogation window). Set the correlation threshold to 0.6.
  12. Click **OK**. After the calculation finishes, save the text file that has the raw data of beads displacement.
  13. Select **Plugins > FTTC > FTTC** and choose the text file in step 9.
  14. Input the **pixel size** (µm), the **Young’s modulus of the gel** (Pascal), and the **plot width and height** based on the experiment and the image of beads.
  15. Click **OK**. The text file that contains the raw data of traction force will be automatically saved in the same directory as the text file in step 9.
2. Utilize Origin to plot the traction field with the same scale for multiple cells (**Figures 1B, 1C, 2B, and 2C**).
  1. Insert the text file that contains the raw data of traction into Excel.
  2. Create a new sheet, input the Y coordinates of traction to the first row (arrange from high values to low values) and the X coordinates to the first column (arrange from low to high).
  3. Input the value of traction to each coordinate from the raw data.
  4. Save the sheet in step 2 as a *.csv file.
  5. Open **Origin** (Version used in our analysis: OriginPro 2017 (Learning Edition)).
  6. Click **File > Open** and import the *.csv file in step 4. Select all the cells and click **Plot > Contour > Contour - Colour Fill**.
  7. In the **Plotting: plotvm** window, select Y across columns. Y values will be automatically set to the first row, and X values will be set to the first column. Next, name the title and click **OK**.
  8. The graph window pops up. Double-click the heatmap.
  9. Click **Levels** in the **Colormap/Contours** window. Next, change the scale level to a reasonable range (0-300 in our analysis) and click **OK**.
  10. Click **Lines**, uncheck **Show on Major Levels Only**, and check **Hide All**. Next, click **OK**.
  11. Right-click the graph and select **Export Graphs…**. Save the image to the specified path.
3. Apply the MATLAB program to calculate the dipole cell traction
  1. Save the traction raw data text file (from step 12 of “Measurement of traction field”) and cell boundary ROI coordinates file (from step 9 of “Measurement of the YAP N/C ratio”) in the same folder.
  2. Open **MATLAB**. Open the folder in step 1 and the dipole traction calculation function file “absdipole.m”.
  3. Read the two text/csv files in step 1 into the MATLAB working space and assign a matrix to two variables (e.g., ‘traction’ and ‘roi’).
  4. Run function ‘absdiple (traction,roi)’.

The first column of the output is the dipole traction force in nN. The second column of the output is the angle of the dipole traction force with respect to the horizontal axis.

## REPRESENTATIVE RESULTS

### 1. Distinct YAP distribution and dynamics in PC9 cancer and B2B normal cells during cell spreading

Representative fluorescent images of YAP distribution in single spreading B2B and PC9 cells on the 5-kPa hydrogel substrate (from the 0^th^ hour to the 10^th^ hour after the cells attached to the substrates) are shown in **Figures 1A** and **2A**, respectively. The B2B cell monotonically increases the spread area over time along with the decrease of the YAP N/C ratio (**Figure 1A**), while the PC9 cell maintains a comparatively unchanging cell spread area, orientation, and YAP N/C ratio throughout the 10-hour spreading process (**Figure 2A**). During the 10-hour of early spreading, the representative B2B cell constitutively deforms the substrate surface and applies time-evolving cell traction across the whole cell area (**Figures 1B and 1C**). In contrast, the representative PC9 cell only develops displacement and traction at the two ends of the cell body and its traction diminishes after 7.5 hours (**Figures 2B and 2C**). Other modes of PC9 cell dynamics are also found (**Figure 6**). In parallel to these different spreading characteristics, B2B and Pc9 cells show distinct YAP distribution and dynamics (**Figure 3**). YAP in B2B cells is concentrated in the nucleus at the 0th hour and becomes more homogeneous distributed across the cell body at the 10th hour. However, PC9 cells show a more homogeneous distribution of YAP in nucleus and cytoplasm throughout the entire 10 hours of the spreading process.

To further investigate the distinct YAP dynamics, we compared the temporal changes in YAP N/C ratio, cell/nucleus area, and traction of multiple single B2B cells (n = 10) and PC9 cells (n = 5) (**Figure 5**). We found that the average YAP N/C ratio of B2B cells decreased from 2.54 ± 0.22 to 1.79 ± 0.21 (n = 10; p-value = 0.0022**; **Figure 5A**) while the average YAP N/C ratio of PC9 cells changed from 1.92 ± 0.26 to 1.57 ± 0.07 (n = 5; p-value = 0.187 (not significant (ns)); **Figure 5A**). The average dipole traction of B2B cells changed from 256.17 ± 123.69 nN to 287.44 ± 99.79 nN (p-value = 0.7593 (ns); **Figure 5B**). The average dipole traction of PC9 cells changed from 141.19 ± 33.62 nN to 168.52 ± 73.01 nN (p-value = 0.7137 (ns); **Figure 5B**). The average cell spread area of B2B cells increased from 613.89 ± 102.43 µm^2^ to 942.51 ± 226.71 µm^2^ (p-value = 0.0512 (ns); **Figure 5C**). The average cell spread area of PC9 cells changed from 495.78 ± 97.04 µm^2^ to 563.95 ± 89.92 µm^2^ (p-value = 0.5804 (ns); **Figure 5C**). The average nucleus spread area of B2B cells increased from 181.55 ± 36.18 µm^2^ to 239.38 ± 43.12 µm^2^ (p-value = 0.1217 (ns); **Figure 5D**) and the average nucleus spread area of PC9 cells changed from 133.31 ± 30.05 µm^2^ to 151.93 ± 22.49 µm^2^ (p-value = 0.5944 (ns); **Figure 5D**). These results suggest that (1) B2B cells show a constitutively higher and more fluctuating YAP N/C ratio over time than that of the PC9 cells; (2) the traction of both B2B cells and PC9 cells show limited variation, while the traction of B2B cells is consistently higher than that of PC9 cells; and (3) in contrast to B2B cells, PC9 cells show limited increase of cell area during the spreading process.

### 2. YAP distribution and dynamics are correlated to the migration states of B2B cells

We compared the YAP N/C ratio and dipole traction of all B2B (n=10) and PC9 (n=5) cells as a function of cell spread area and nucleus spread area. The YAP N/C ratio and dipole traction of PC9 cells do not show a clear correlative trend within their small cell and nucleus spread area ranges (**Figure 6**). In contrast, the YAP N/C ratio and dipole traction of B2B cells appear to follow two distinct trends (**Figures 6A and 6C**), suggesting there might be two groups of B2B cells that co-exist in our experiment. In the first group, the YAP N/C ratio and dipole traction increase along with the enlargement of the cell spread area and reach their maximums at ∼ 1000 µm^2^ (**Figure 6C and 6D**, indicated by the yellow dashed line). In the second group, the YAP N/C ratio and dipole traction increase at a slower rate with the enlargement of the cell spread area and maintain nearly constant when the cell spread area continues to increase (**Figures 6C and 6D**, indicated by the green dashed line).

### 3. PC9 cancer cells generate tractions in peri-nuclear regions

For single spreading PC9 cells, we found they evidently displace the substrates at the peri-nuclear regions, starting from the 6^th^ hour of culture on. To visualize the peri-nuclear displacement caused by cell traction, we overlapped the images of fluorescent beads taken before (red) and after (green) the removal of the cells from the substrates (please see the Protocol section for details). The beads that do not have any displacement will appear yellow in the overlapped images, i.e., the addition of red and green colors, while the beads that displace apart from their resting positions due to the cell traction will show separated green and red colors. Noticeably, in both PC9 (**Figures 7C and 7D**) and B2B (**Figure 7E**) cells, we observed beads displacement in the cytoplasm and within the nucleus, in addition to those at the cell boundary. To highlight the peri-nuclear displacement, we use the Boussinesq equation to predict the 2D theoretical displacement generated by a hypothetical dipole force at the cell boundary (black dashed line in **Figure 7B**) [13]. Comparing this theoretical curve with the real substrate displacement measured along the same axis (white dashed line in **Figure 7D**), we found that the real displacements within the nucleus are indeed 1.5–8-fold-larger than the theoretical value (**Figure 7B**), indicating the existence of traction force at the peri-nuclear regions.

## DISCUSSION

Mechanotransducers YAP/TAZ are new promising therapeutic targets for the development of cancer therapies [14]–[16]. Emerging data suggest that YAP/TAZ promote the proliferation and invasion of cancer cells. Mechanics-induced YAP translocations from cytoplasm to nucleus activate transcriptions of genes related to cell migration, proliferation, invasion, and apoptosis, leading to aberrant cell behaviors [17]–[20]. In this research, we aim to explore the potential correlation of the YAP N/C ratio and cell mechanics in two typical human cancer and normal cell lines.

We found that the YAP N/C ratio of PC9 cancer cells is constitutively lower than that of B2B normal cells (**Figure 5A**). This relationship is contradictory with the majority of published findings that compare YAP concentration in nucleus between normal and cancer cells. Most findings show that YAP is more concentrated in the nucleus of cancer cells than that in normal cells [16], [17]. Only one study reported an exception in breast cancer research [21]. To the best of our knowledge, our data is the first one to show a lower YAP N/C ratio in a human lung cancer cell line. We hypothesize that the reason for a stable YAP N/C ratio in PC9 cells might relate to the fact that the cell/nucleus spread area and traction in PC9 cells have little variation at the early spreading stage. The dissection of underpinning molecular mechanisms of low YAP N/C ratio in PC9 and B2B cells is ongoing in our laboratories.

During the first 10 hours of spreading, these two cell lines show a distinct relationship between the YAP N/C ratio, cell traction, and spread areas (**Figure 5**). For B2B cells, a higher YAP N/C ratio is correlated with the higher cell and nucleus spread area (**Figures 6A and 6B**), which is consistent with the reported data of other normal cells [22]. Interestingly, although the developmental trend of this relationship is generally found in all B2B cells recorded, we found two different degrees (high and low) of this relationship. For B2B cells that spread and migrate simultaneously, they show lower traction and higher cell and nucleus spread area with higher YAP N/C ratio (2.05 ± 0.32). For B2B cells that spread and remain at the same location, they show higher traction and lower cell and nucleus spread area with lower YAP N/C ratio (1.74 ± 0.21). These two degrees of relationships are demonstrated in the bifurcated scattered data groups (**Figures 6C and 6D**). As reported in literature, stationary normal cells, such as embryonic fibroblast NIH 3T3 cell, have higher traction compared to that of migratory cells [23]. Our data suggest that the spreading and non-migrating B2B cells applied higher traction than that of spreading and migrating B2B cells, likely suggesting that high traction is needed for non-migrating cells to stabilize on the substrate.

In addition, our data show that stationary normal B2B cells generate higher peri-nuclear force, while previous research conducted by other groups reported only higher cell periphery traction in stationary cells [23]–[26]. We think that the difference in the intrinsic tendency of migration in the experiments might cause these contradictory results. Square-shape micropatterning had been used in the published experiments to confine single cells spreading and inhibit migration but whether the cells have the tendency to migrate is unknown. Since migratory cells show high traction force at the periphery of the cells [27], chances are that cells with the tendency to migrate will still maintain high periphery traction even their migration is restricted. In our results, the stationary cells are not restricted by any micropattern and thus do not tend to migrate. Another possibility is that the cell shape defined by micropattern may affect the distribution of focal adhesion and traction force [28]. Our result is generated without any confining micropatterning and represents the force distribution of stationary cells at their original shape.

To the best of our knowledge, only one publication to date specifically reported the finding of peri-nuclear forces in normal cells (mouse embryonic fibroblasts) potentially caused by actin-cap spanning across the nucleus [29]. YAP cytoplasm-to-nucleus translocation correlates with the increase of peri-nuclear force [29]. We do not find any publications available that report peri-nuclear force or actin-cap in cancer cells. An indirect study on melanoma cancer cells demonstrated that the actin rim (another peri-nuclear actin organization that is located around but not covering the nucleus) reduces cell migration rates [30], indirectly suggesting the existence of peri-nuclear force. However, no direct experimental data are reported. In our research, we found both PC9 and B2B cells show peri-nuclear displacement and traction. The mechanisms of the generation of the peri-nuclear forces and their effects remain controversial. In normal cells, actin-cap was reported to have a role in regulating nucleus morphology and chromatin organization [31], transmit mechanical signals from focal adhesion into nucleus through linkers of nucleoskeleton and cytoskeleton (LINC) complex [32], and regulate cell migration [33]. Lamin A/C is related to the formation and disruption of actin-cap [29]–[33]. However, the report that claimed actin-cap generates the peri-nuclear force did not consider the potential role of actin-rim[29]. In cancer cells, overexpression of Lamin A facilitates the formation of actin-rim and restricts the cancer cell migration. Overexpression of Lamin B reduces the actin-rim formation and promotes migration[30]. Peri-nuclear force might be involved in this process due to the existence of peri-nuclear actin organization and the effect of Lamin A. However, in this research, no data show the measured peri-nuclear forces or the behavior of actin-cap. Therefore, our discovery of peri-nuclear force in PC9 cells is the first report showing peri-nuclear forces and displacements in lung cancer cells. We are currently investigating the molecular mechanisms and functions of peri-nuclear forces, using both PC89 and B2B cells as the test beds.

Beyond the all-optical mechanobiology interrogation that is demonstrated in this paper, our integrated multi-functional system can be applied to optically probe a myriad of other physiological and pathobiological signals in living systems. For example, our laboratory has recently established multiple stably transduced human cancer cell lines that co-express three light-responsive membrane proteins: membrane voltage indicator QuasAr2 (excitation: 640 nm; emission: 660 nm–740 nm), membrane voltage depolarizer CheRiff (excitation: 488 nm), and membrane voltage hyperpolarizer eNpHR3 (excitation: 590 nm) (**Figures 8A, 8B and 8C**). These three functional proteins can be activated by spectrum-orthogonal laser lines in a crosstalk-free manner, enabling all-optical two-way signaling communications (readout and control) of membrane electrophysiology. This is a new version of “Optopatch (optical patch-clamp)”. Using our integrated system and a manual patch-clamp (**Figure 8F**), we have validated the all-optical control and readout of the membrane voltage (*V*_*m*_) in single human cancer cells and cell spheroids (**Figures 8D and 8E**). The new all-optical electrophysiology (Optopatch) interrogation opens the possibility for detailed explorations of previously inaccessible bioelectricity in cancer cells, which potentially may advance our knowledge of tumor biology from a new axis.

## ACKNOWLEDGMENTS

This project is financially supported by the Cancer Pilot Award from UF Health Cancer Center (X. T. and D. S.) and the Gatorade Award Start-up Package (X. T.). We sincerely appreciate the intellectual discussions with and the technical supports from Dr. Jonathan Licht (UFHCC), Dr. Rolf Renne (UFHCC), Dr. Ji-Hyun Lee (Biostatistics, UF), Dr. Hugh Fan (MAE, UF), Dr. Warren Dixon (MAE, UF), Dr. Ghatu Subhash (MAE, UF), Dr. Malisa Sarntinoranont (MAE, UF), Dr. Scott Banks (MAE, UF), Dr. Matthew Traum (MAE, UF), Dr. Dr. David Hahn (University of Arizona), Dr. Weihong Wang (Oracle Corporation), Dr. Youhua Tan (Hong Kong Polytechnic University), and Support Team of Nikon (Drs. Jose Serrano-Velez, Larry Kordon, and Jon Ekman). We are deeply grateful for the generous and effective supports from all members of Tang’s, Siemann’s, and Guan’s research laboratories and all staff members of the MAE & ECE Departments, UF.

## References

[1] C. Werley, S. Boccardo, A. Rigamonti, E. Hansson, and A. Cohen, “Multiplexed Optical Sensors in Arrayed Islands of Cells for multimodal recordings of cellular physiology,” Nat. Commun., vol. 11, no. 1, pp. 1–17, 2020.

[2] B. Yang et al., “Epi-illumination SPIM for volumetric imaging with high spatial-temporal resolution,” Nat. Methods, vol. 16, no. 6, pp. 501–504, 2019.

[3] A. Saraswathibhatla, E. E. Galles, and J. Notbohm, “Spatiotemporal force and motion in collective cell migration,” Sci. Data, vol. 7, no. 1, pp. 1–7, 2020.

[4] A. Saraswathibhatla, S. Henkes, E. E. Galles, R. Sknepnek, and J. Notbohm, “Coordinated tractions control the size of a collectively moving pack in a cell monolayer,” arXiv preprint 2102.09036, 2021.

[5] Q. Luo et al., “Automatic Multi-functional Integration Program (AMFIP) towards All-optical Mechanobiology Interrogation,” BioRxiv, 2021.

[6] A. Edelstein, N. Amodaj, K. Hoover, R. Vale, and N. Stuurman, “Computer control of microscopes using manager,” Curr. Protoc. Mol. Biol., vol. 92, no. 1, pp. 14–20, 2010.

[7] A. Tulpule et al., “Cytoplasmic protein granules organize kinase-mediated RAS signaling,” BioRxiv, p. 704312, 2019.

[8] S. Feng, S. Sekine, V. Pessino, H. Li, M. D. Leonetti, and B. Huang, “Improved split fluorescent proteins for endogenous protein labeling,” Nat. Commun., vol. 8, no. 1, pp. 1– 11, 2017.

[9] X. Tang, A. Tofangchi, S. Anand, and T. Saif, “A novel cell traction force microscopy to study multi-cellular system,” PLOS Comput. Biol., vol. 10, no. 6, p. e1003631, 2014.

[10] X. Tang et al., “Mechanical force affects expression of an in vitro metastasis-like phenotype in HCT-8 cells,” Biophys. J., vol. 99, no. 8, pp. 2460–2469, 2010.

[11] J. Schindelin et al., “Fiji: An open-source platform for biological-image analysis,” Nat. Methods, vol. 9, no. 7, pp. 676–682, 2012.

[12] J. L. Martiel et al., “Measurement of cell traction forces with ImageJ,” Methods Cell Biol., vol. 125, pp. 269–287, 2015.

[13] I. A. Okumurai, “On the generalization of Cerruti’s problem in an elastic half-space,” Doboku Gakkai Ronbunshu, vol. 1995, no. 519, pp. 1–10, 1995.

[14] S. Piccolo, S. Dupont, and M. Cordenonsi, “The biology of YAP/TAZ: hippo signaling and beyond,” Physiol. Rev., vol. 94, no. 4, pp. 1287–1312, 2014.

[15] W. Hong and K. L. Guan, “The YAP and TAZ transcription co-activators: Key downstream effectors of the mammalian Hippo pathway,” Semin. Cell Dev. Biol., vol. 23, no. 7, pp. 785– 793, 2012.

[16] F. Zanconato, M. Cordenonsi, and S. Piccolo, “YAP/TAZ at the Roots of Cancer,” Cancer Cell, vol. 29, no. 6, pp. 783–803, 2016.

[17] Y. Wang, Q. Dong, Q. Zhang, Z. Li, E. Wang, and X. Qiu, “Overexpression of yes - associated protein contributes to progression and poor prognosis of non-small-cell lung cancer,” Cancer Sci., vol. 101, no. 5, pp. 1279–1285, 2010.

[18] H. Li et al., “Inhibition of YAP suppresses CML cell proliferation and enhances efficacy of imatinib in vitro and in vivo,” J. Exp. Clin. Cancer Res., vol. 35, no. 1, pp. 1–11, 2016.

[19] X. Tang et al., “A mechanically-induced colon cancer cell population shows increased metastatic potential,” Mol. Cancer, vol. 13, no. 1, pp. 1–15, 2014.

[20] T. Panciera, L. Azzolin, M. Cordenonsi, and S. Piccolo, “Mechanobiology of YAP and TAZ in physiology and disease,” Nat. Rev. Mol. Cell Biol., vol. 18, no. 12, pp. 758–770, 2017.

[21] M. Yuan et al., “Yes-associated protein (YAP) functions as a tumor suppressor in breast,” Cell Death Differ., vol. 15, no. 11, pp. 1752–1759, 2008.

[22] N. Koushki et al., “Lamin A redistribution mediated by nuclear deformation determines dynamic localization of YAP,” BioRxiv, 2020.

[23] S. S. Chang, A. D. Rape, S. A. Wong, W. hui Guo, and Y. li Wang, “Migration regulates cellular mechanical states,” Mol. Biol. Cell, vol. 30, no. 26, pp. 3104–3111, 2019.

[24] J. Lee, A. Abdeen, X. Tang, T. Saif, and K. Kilian, “Geometric guidance of integrin mediated traction stress during stem cell differentiation,” Biomaterials, vol. 69, pp. 174– 183, 2015.

[25] J. Lee, A. Abdeen, X. Tang, T. Saif, and K. Kilian, “Matrix directed adipogenesis and neurogenesis of mesenchymal stem cells derived from adipose tissue and bone marrow,” Acta Biomater., vol. 42, pp. 46–55, 2016.

[26] X. Tang, P. Bajaj, R. Bashir, and T. Saif, “How far cardiac cells can see each other mechanically,” Soft Matter, vol. 7, no. 13, pp. 6151–6158, 2011.

[27] M. Dembo and Y. L. Wang, “Stresses at the cell-to-substrate interface during locomotion of fibroblasts,” Biophys. J., vol. 76, no. 4, pp. 2307–2316, 1999.

[28] Y. W. Andrew D. Rape, Wei-hui Guo, “The regulation of traction force in relation to cell shape and focal adhesions,” Biomaterials, vol. 32, no. 8, pp. 2043–2051, 2011.

[29] J. Y. Shiu, L. Aires, Z. Lin, and V. Vogel, “Nanopillar force measurements reveal actin-cap-mediated YAP mechanotransduction,” Nat. Cell Biol., vol. 20, no. 3, pp. 262–271, 2018.

[30] A. Fracchia, T. Asraf, M. Salmon-Divon, and G. Gerlitz, “Increased Lamin B1 Levels Promote Cell Migration by Altering Perinuclear Actin Organization,” Cells, vol. 9, no. 10, p. 2161, 2020.

[31] N. M. Ramdas and G. V. Shivashankar, “Cytoskeletal control of nuclear morphology and chromatin o1rganization,” J. Mol. Biol., vol. 427, no. 3, pp. 695–706, 2015.

[32] S. B. Khatau et al., “A perinuclear actin cap regulates nuclear shape,” Proc. Natl. Acad. Sci. U. S. A., vol. 106, no. 45, pp. 19017–19022, 2009.

[33] D. H. Kim, S. Cho, and D. Wirtz, “Tight coupling between nucleus and cell migration through the perinuclear actin cap,” J. Cell Sci., vol. 127, no. 11, pp. 2528–2541, 2014.

